# Neural correlates of age-related changes in social decisions from episodic memory

**DOI:** 10.1101/2024.08.26.609251

**Authors:** Camilla van Geen, Michael S. Cohen, Karolina M. Lempert, Kameron A. MacNear, Frances M. Reckers, Laura Zaneski, David A. Wolk, Joseph W. Kable

## Abstract

Older adults are frequent victims of financial scams. Previous behavioral research suggests that this may be due to systematic biases in how they make decisions about whom to trust: for instance, Lempert et al. (2022) found that relative to younger adults, older adults were more likely to base decisions about whether to re-engage with someone on how generous that person looked, rather than on their memory for how they had previously behaved. Here, we aimed to identify the neural correlates of these age-dependent changes in social decision-making in order to clarify the mechanism by which they emerge. Using functional Magnetic Resonance Imaging (fMRI), we measured neural activity while a total of 86 participants – 45 younger and 41 older adults – learned about how much of a $10 endowment an individual, represented by a picture of their face, was willing to share with them in a dictator game. After this encoding phase, participants then made decisions about whom they wanted to play another round of the dictator game with. In line with previous findings, we found that older adults did not reliably prefer to re-engage with people who had proven themselves to be generous. This bias was the result of several factors: (1) older adults had worse associative memory for how much each person had shared, possibly due to an age-dependent decrease in neural activity in the medial temporal lobe (MTL) during encoding, (2) older adults had a stronger tendency to re-engage with familiar over novel faces regardless of their past behavior, and (3) while activity in value-responsive brain regions tracked with how generous a face looked across the age range, older adults were less able to inhibit the influence of these irrelevant perceptual features when it was necessary to do so. In line with this behavioral effect, younger adults showed greater activation in the inferior frontal gyrus (IFG) during choices that required suppressing irrelevant perceptual features in favor of associative memory. Taken together, our findings highlight age-dependent changes in both the ability to encode relevant information and to adaptively deploy it in service of social decisions.

## Introduction

As life expectancy rises, older adults are projected to make up a growing portion of the population: by certain estimates, 23% of Americans are expected to be older than 65 in 2050 (U.S. Census Bureau). Older adults also hold a considerable amount of wealth and tend to be over-represented in positions of power. Despite this substantial and growing influence, however, older adults are also disproportionately victimized by financial scams. In 2020, for instance, 28% of the $4.1 billion stolen through scams came from people over the age of 65 (2020 Elder Fraud Report, FBI). Recent studies suggest that one of the reasons for this phenomenon is that older adults display systematic biases in how they make decisions about whom to trust (Bailey & Leon, 2019; Lempert et al., 2022; Rosi et al., 2019). Here we aim to identify the neural correlates of these age-dependent changes in social decision-making, leveraging what is known about the aging brain to better understand the mechanisms by which these changes emerge.

When people choose whether to interact with someone, they bring multiple sources of information to bear on their decisions. Two particularly influential factors are: (1) judgments based on superficial attributes like facial features, and (2) memory for how people behaved during previous interactions. Indeed, we form impressions of others based on their appearance, with certain physical traits leading to perceptions of higher approachability (Bar et al., 2006; Todorov et al., 2008). These stereotypes do not necessarily track reality, but they nonetheless modulate approach behavior and impact subsequent memory (Cassidy et al., 2012; Zebrowitz & Montepare, 2008). For instance, individuals who look more “baby-faced” are reliably perceived as more approachable and less dominant, even in how they are subsequently remembered after conflicting information is provided (Berry & McArthur, 1986; Cassidy et al., 2012).

Social decisions also often involve repeated interactions with the same person – in these cases, memory for previous experiences can be recruited to decide whether to re-engage. Indeed, memory serves a fairly ubiquitous role in decision-making, providing crucial information about the value of available options (Biderman et al., 2020; Schlichting & Preston, 2015; Shadlen & Shohamy, 2016; Zeithamova et al., 2012). Historically, research on mnemonic contributions to decision-making has focused on how we learn value from repeated experiences (for example, in reinforcement learning – Schönberg et al., 2007; Sutton & Barto, 1998). However, people can also encode the value of an option after a single experience and retrieve it when necessary. Several recent papers have shown that young adult participants learn the value of trial-unique images and then leverage this information to subsequently make adaptive choices (FeldmanHall et al., 2021a; Lempert et al., 2022b; Murty et al., 2016; Nicholas et al., 2022).

Both of these influences on social decisions – perceptually-based stereotypes and memory for previous interactions – can be affected by aging. On one hand, older adults exhibit deficits in inhibitory control, especially when it comes to disregarding irrelevant semantic and perceptual information (Amer et al., 2016, 2022; Dey et al., 2017; Healey et al., 2013; Yoss et al., 1970). In the context of social decision-making, older adults may be more susceptible to stereotypes, especially when a person looks approachable. Previous research on social cognition has identified a “positivity bias” that emerges with age, such that older adults prioritize positively-valenced social information in both choice and memory (Carstensen & Mikels, 2005; Mather & Carstensen, 2005). On the other hand, older adults also exhibit deficits in episodic memory, especially when it requires combining multiple features – like value and stimulus identity (Hoyer & Verhaeghen, 2006; Naveh-Benjamin et al., 2003, 2004; Old & Naveh-Benjamin, 2008; Wolk et al., 2013). On the neural level, aging is associated with a marked decrease in the volume of the hippocampus (Driscoll et al., 2003; O’Shea et al., 2016), and the degree of this decrease has been linked to poorer memory when association of features is required (Wolk et al., 2011). Taking these sets of findings together, we might expect that aging decreases the contribution of episodic memory to decision-making, leaving room for a greater reliance on irrelevant perceptual features like social stereotypes.

In a recent behavioral study, Lempert et al. (2022) found evidence that older adults do indeed over-rely on perceptual features to guide social choices while simultaneously having worse memory for previous interactions. In that study, which used a similar task design as the present work, participants received one-shot information about how much a person (denoted by a picture of their face) had chosen to share with them in a dictator game (Güth et al., 1982). Participants then made decisions about whom they would prefer to play another round of the game with, knowing that anyone who had previously been generous would continue to be, and vice versa (Lempert et al., 2022). Importantly, the faces varied in how generous they looked, but there was no relationship between true and perceived generosity. As such, adaptive behavior consisted of choosing to play with previously rewarding partners, and avoiding those who did not share. In this study, memory for how much each person had shared declined with age, as did accurate value-based decisions. Specifically, older adults were more biased towards misremembering faces as generous, tended to approach familiar faces regardless of their previous behavior, and were more likely to make choices influenced by irrelevant facial features.

Although this behavioral study reveals interesting biases in how older adults approach social decisions, several important questions remain about how these biases emerge. For instance, it is unclear whether older adults’ memory deficits are due to disrupted processing at encoding or at retrieval. Furthermore, we do not know whether older adults are more likely to base their decisions on stereotypical facial features because they are more sensitive to stereotypes than young adults, or rather because they have more difficulty inhibiting the influence of these stereotypes at the time of choice. Neuroimaging data can help address these outstanding questions by localizing age-related differences during encoding, retrieval, and choice to specific constructs, brain regions, and moments in time.

To this end, we used functional Magnetic Resonance Imaging (fMRI) to measure brain activity while a total of 86 younger and older adults participated in the memory-based decision-making task described in Lempert et al (2022). Participants first learned about how much of a $10 endowment an individual, represented by a picture of their face, was willing to share with them. Later, they were presented with these faces again, and made decisions about with whom to play another round. We found an age-related decrease in neural activity in the medial temporal lobe (MTL) during encoding, which was correlated with worse decision-making performance. Furthermore, although both younger and older adults were sensitive to facial features – as indexed both neurally and behaviorally – older adults were less able to adaptively inhibit the influence of these features on choice. Our findings aim to disentangle the complex forces that shape social decision-making and highlight age-dependent changes in both the ability to encode relevant information and to adaptively deploy it.

## Methods

### Participants

A total of 86 participants completed the study: 41 older adults (mean age 72.7 years; range 61-86 years, 20 male, 21 female) and 45 younger adults (mean age 26.5 years; range 21-37 years, 20 male, 25 female). The older adults were recruited from the Clinical Core of the University of Pennsylvania’s Alzheimer’s Disease Research Center (ADRC). As part of their involvement in the center, participants undergo annual psychometric testing as defined by the Uniform Data Set 3.0 (UDS) and medical/neurological examination. Older adults who participated in this study were all designated as cognitively normal based on this assessment and consensus conference determination. The young adult sample was recruited through the Penn MindCORE online participant recruitment system. All participants provided informed consent, and the procedures were approved by the University of Pennsylvania’s Institutional Review Board.

### Procedure

In this study, we used the same experimental stimuli, reward task, decision task, and memory task as Lempert et al. (2022). The main difference is that in this sample, participants completed the experiment while undergoing fMRI scanning. Although half of the trials involved non-social stimuli (houses), we focus our analyses here on decisions involving social stimuli (faces). This is because Lempert et al. found several effects of age that specifically impacted social decisions, such that even when older adults had intact memory for how much a specific person had given them previously, they did not necessarily choose adaptively. Our analyses focus on identifying neural activity associated with the effects of age on memory, social biases, and their interaction.

### Stimuli

Prior to this study, a separate group of participants was invited to the lab to complete decision-making questionnaires and to have their photograph taken. They were told that they were contributing to a database of decisions and images that could be used for future studies. One of the decision-making tasks they completed was a standard dictator game: participants had to decide how much of a $10 allotment they would share with an anonymous other. The available options varied from $0 to $10, in one-dollar increments. To create our experimental stimuli, we selected a subset of these participants, composed of people who chose either an even split ($5/$5) or to keep all of the money for themselves ($10/$0). We then used the photographs of their faces as stimuli for the reward, decision, and memory tasks, selecting an equal number of people who had given $5 and $0. In order to account for potential race or gender-related confounds, we ensured representation across these characteristics. Our stimulus set of faces in the reward task contained four Asian females, four Asian males, four Black females, four Black males, eight White females, and eight White males. Individuals in each of these sub-groups were evenly split between the high value ($5/$5) and low value ($10/$0) categories.

Once the stimulus set was created, we then invited a separate group of independent raters (*n* = 20) to evaluate the faces on a variety of characteristics including attractiveness, trustworthiness, dominance, and competence. We also asked raters how much they thought each person would share with them in a hypothetical dictator game, from $0 to $10 (see Figure 1A). This variable, which we refer to as perceived generosity, is what we will use in subsequent analyses as a measure for how generous each face looked. As described in Lempert et al. (2022), there was no correlation between perceived generosity and actual generosity. The absence of any correlation simplifies our analyses, since it allows us to decouple choices based on perceived generosity from choices based on episodic memory.

**Figure 1.**
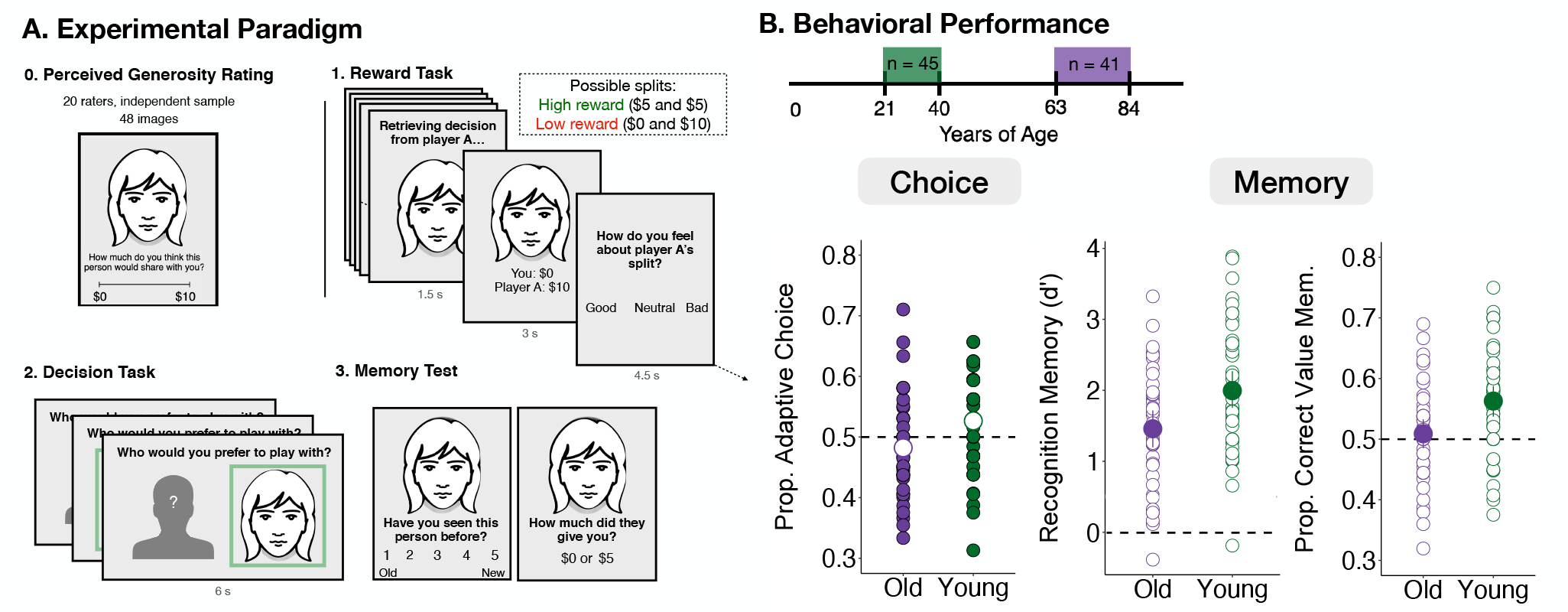
**(A)** Overview of the experimental paradigm. In the reward task, participants see 32 faces and learn how much each person shared with them in a dictator game (in the actual task, images are photographs of real people’s faces, which we omit here to protect their privacy). 16 people shared $0 (low reward) and 16 people shared $5 (high reward). During the decision task, participants are asked whether they would prefer to play another round of the dictator game against a known opponent (face) or someone at random (outline). All of the faces from the reward task are shown again, as well as 16 novel faces. Finally, the memory test probes participants’ recognition memory for faces from the previous tasks, as well as their associated value. **(B)** Choice and memory accuracy across age groups. As a group, young adults tend to choose to play again with high value faces and avoid low value faces, while older adults are not above chance. In the memory phase, both younger and older adults can identify which faces they have seen before. Only younger adults are above chance in their ability to remember the associated value, however.

### Reward Task

On each trial of the reward task, participants were first shown a trial-unique image of a face accompanied by the text, “Retrieving decision from Player A” (see Figure 1A). Participants knew that Player A, whose face was displayed, had decided how much of a $10 allotment to share with them. After 1.5 seconds, the text was replaced to reveal the outcome of the trial, which was always either $5/$5 (high reward) or $0/$10 (low reward). The outcome remained on the screen for 3 seconds, after which participants had 4.5 seconds to rate how they felt about the split (good, neutral, or bad). After a jittered intertrial interval between 1 and 5 seconds (drawn from a normal distribution with a mean of 3s ± 1), the next trial began with the next Player A. In total, participants saw 32 trial-unique faces, equally split between high and low reward. The order of trials was randomly generated prior to data collection but held constant across participants. For our fMRI analyses, we model the entire time the face was on the screen as a single event, combining the first 1.5 and 3 seconds into one boxcar regressor.

### Distractor Task

In between the reward and decision tasks, participants completed a 5-minute anagrams task in which they were presented with a series of scrambled words and given 30 seconds to unscramble each one. Although participants remained in the scanner for the duration of this distractor task, no MRI data were collected during this period; the microphone was left on, however, so that we could ensure that participants were attempting each anagram. The task automatically moved on to the next trial after 30 seconds on a given word.

### Decision Task

On each trial of the decision task, participants chose which of two people they would rather play another round of the dictator game with. One person’s face was always presented on the right side of the screen, while a schematic outline of an indeterminate person was shown on the left. Participants were told that choosing the schematic meant a Player A would be chosen at random from the database, with equal probability that they would share $0 or $5. For 32 out of the 48 decision trials, the person’s face on the right was one that participants had learned about during the reward task, meaning that they could use that information to guide their choice. For the other 16 trials, the face was novel, so participants could not use value memory to inform these decisions. On each trial, participants had 5 seconds to make a decision. After making their choice, a box appeared on the screen for one second, indicating which option the participant chose. For the fMRI analyses, we model both the choice (5s) and feedback phase (1s) as a single event, totaling 6 seconds. After feedback, a fixation cross was displayed on screen for a randomly determined amount of time between 1 and 5 seconds before the next decision trial. At the end of the study, we paid out one of the participants’ decisions at random, ensuring that their choices throughout the task were incentive compatible. If the randomly selected trial was one in which the participant had chosen the face, they received what that person offered in the Dictator Game. If the randomly selected trial was one in which the participant chose the schematic, we drew $0 or $5 with 50% probability.

### Memory Task

After participants completed the reward and decision tasks, they exited the scanner before taking a surprise memory test. During the memory test, participants saw all the faces from the reward phase, interleaved with 16 novel faces. Trial order was randomized before the start of data collection, but then held constant across participants. To assess participants’ recognition memory, we first asked them to report on a scale from 1 (highly confident yes) to 5 (highly confident no) whether that they had seen a given face during the reward task. If participants responded 1, 2, or 3 during this initial test of recognition memory, they then received a value memory prompt, which asked whether the face was associated with $0 or $5. They were also asked to indicate their confidence in this value memory from 1 (just guessing) to 3 (very confident). Finally, we asked participants whether they had chosen to play with that particular person during the decision task. We allowed participants to take as long as they needed to make their responses, with a one-second interval between each trial.

### Behavioral Data Analysis

All behavioral analyses were conducted in R (version 4.2.3).

#### Decision-Making

To measure whether age affected overall performance in the decision task, we first compared the average proportion of adaptive choices across groups (Figure 1B). To do so, we coded a decision as adaptive if participants chose a high value face or avoided a low value face. We calculated each participant’s percentage of adaptive choices and conducted a one-sample *t*-test for each group to assess whether performance was reliably different from chance (corresponding to 0.5, since two possible choices exist). We also ran an independent samples *t*-test on participants’ accuracy scores in order to assess whether age group significantly modulated task performance.

Next, to assess whether participants used memory to make their choices, we used the “lme4” package in R (Bates et al., 2013) and ran a multi-level logistic regression estimating the probability that a participant chose the person as a function of the interaction between how much that person shared during the reward task ($0 or $5) and the participant’s value memory accuracy (0 or 1). To capture age effects, we also ran a version of this regression only including trials where participants had intact value memory. In this regression, we included the interaction between reward and age group to predict choice, as well as a random intercept for each participant.

To quantify the influence of perceived generosity on choice, we again followed a regression-based approach. As the main predictor of interest, we used the z-scored perceived generosity rating for each face. We allowed the influence of perceived generosity to vary for each participant, along with the intercept. We also included age group as both a main effect and an interaction term. In our analysis that focuses on the effect of perceived generosity specifically on low-value trials, we ran a multi-level logistic regression looking at the effect of perceived generosity on choice only on trials in which low-value faces were presented. We allowed the main effect to vary for each participant and then assessed age differences by running a two-sample *t*-test on the random effect coefficients for the effect of perceived generosity on choice.

Finally, we assessed participants’ tendency to choose faces over the schematic by computing the proportion of trials on which participants chose the face, and then running a one-sample *t*-test compared to 0.5. If the proportion is reliably above 0.5, this suggests the presence of a face bias. To measure age effects, we also conducted a two-sample *t*-test comparing the proportion of trials on which younger versus older adults chose the face.

#### Memory

To analyze recognition memory, we used a signal detection theory approach. First, we excluded memory trials where participants said they were guessing (responded 3 on the 1-5 scale). On average, we excluded 5.2 trials per participant (± 4.1). We then computed the number of hits (old images marked as old) and false alarms (new images marked as old) for each participant. The z-scored difference between these two metrics corresponds to d’, which we take as our primary measure of recognition memory performance. d’ measures the extent to which each participant can reliably tell apart old images from new ones. In order to assess whether participants’ recognition memory was above chance, we ran a one-sample *t*-test on each group’s performance, comparing the distribution of d’ to 0. We also subjected the d’ scores to an independent-samples *t*-test in order to test for age effects.

To measure value memory performance, we calculated the proportion of times participants reported correct value memory, out of the total number of hits. We then used *t*-tests to measure whether this metric was reliably above chance (0.5), as well as whether it was significantly different across age groups. We also analyzed value memory using the signal detection approach described in Lempert et al. (2022). Namely, we considering an item a *hit* if a $5 face was remembered as such, a *correct rejection* if a $0 face was remembered as such, a *false alarm* if a $0 face was remembered as $5, and a *miss* if a $5 face was remembered as $0. This allowed us to calculate an “associative memory bias” reflecting the tendency to remember faces as having shared $5 rather than $0.

### fMRI Methods

#### Data Acquisition and Preprocessing

Magnetic resonance images were acquired with a 3T Siemens Magnetom Prisma scanner and a 64-channel head coil. A three-dimensional, high-resolution structural image was acquired using a T1-weighted, magnetization-prepared, rapid-acquisition gradient-echo pulse sequence (voxel size=0.8×0.8×0.8mm; matrix size=241×286; 241 axial slices; repetition time=3000 ms; echo time= 30 ms). While participants completed the task, functional images were acquired using a T2*-weighted gradient echo-planar imaging pulse sequence (voxel size=2×2×2 mm; interslice gap=0.15 mm; matrix size=98×98; 72 oblique axial slices; repetition time=1500 msec; echo time= 30 ms). Slices were angled +30° with respect to the anterior commissure–posterior commissure line to reduce signal dropout in the OFC (Weiskopf et al., 2006).

We preprocessed the data using *fMRIPrep* 20.2.2 (Esteban et al., 2019), which is based on *Nipype* 1.6.1 (Gorgolewski et al., 2011). All BOLD runs were motion corrected, slice-time corrected, b0-map unwarped, registered, and resampled to a Montreal Neurological Institute (MNI) 2-mm template. After preprocessing, we performed spatial smoothing in FSL using a Gaussian kernel with a full-width half maximum (FWHM) of 5mm.

#### fMRI Analyses

We conducted all univariate fMRI analyses in FSL, using generalized linear models (GLMs) to measure changes in neural activity in response to our conditions of interest. For each model, we also included confound regressors to account for motion-related artifacts. These consisted of 6 realignment parameters (three translational and three rotational), as well as the temporal derivative, quadratic terms, and the quadratic of the temporal derivative of each. Additionally, we performed spike regression by creating as many unit impulse functions are there were volumes with framewise displacement > 0.9 mm for each participant. Each of these regressors had a value of 1 at the timepoint where framewise displacement was larger than 0.9mm, and a value of 0 elsewhere (Ciric et al., 2017). The number of spike-regressors varied substantially across participants, ranging from 0 to 82 with a median of 2. Group-level maps were computed using a FLAME1 + FLAME2 mixed effects model, thresholded at *z* = 3.1, and corrected for multiple comparisons using parametric cluster-based correction.

We used a recently developed python-based pipeline (https://github.com/alicexue/fmri-pipeline) to help automate the analyses in FSL. As described in the results section, we focused on the 4.5 seconds during which each face was on the screen for the reward task analyses, and the 6 seconds of decision + feedback time for the decision task analyses. To measure differences in neural activity during reward encoding (Figure 2), we included the following regressors of interest in our GLM: 1) high value faces; 2) low value faces, 3) parametric modulator for response time, 4) parametric effect of perceived amount given for all faces. To examine age effects at encoding, we created a contrast that combined regressors for high and low value faces and examined the effect of age group at the group-level. Figure 2A illustrates voxels where activity was significantly modulated by age for this contrast, and supplementary table 1 lists the coordinates of each significant cluster.

**Figure 2.**
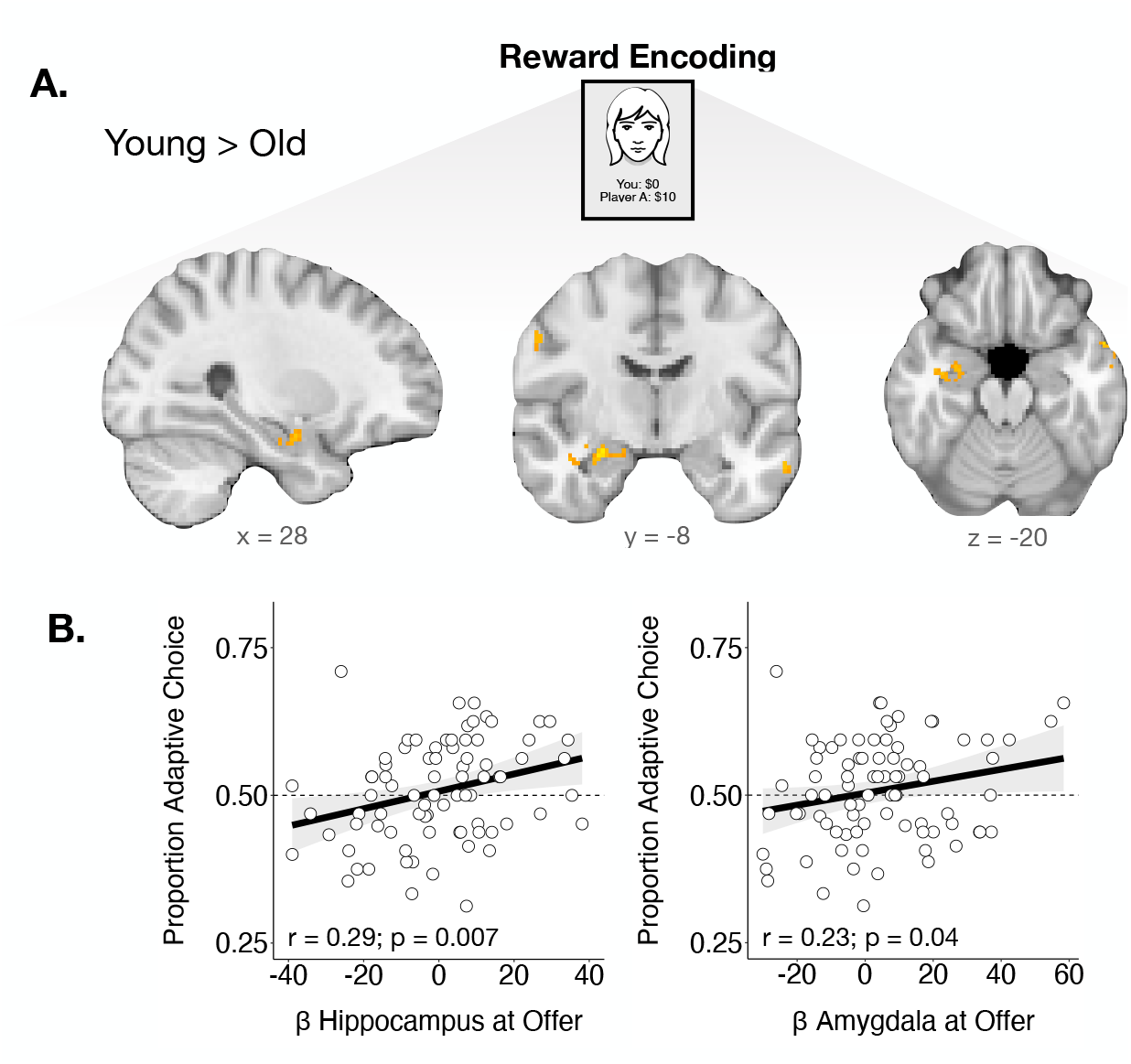
**(A)** Age differences in neural activation during reward encoding. Relative to older adults, young adults exhibit more neural activity in the amygdala and anterior portion of the hippocampus. **(B)** Relationship between choice accuracy in the decision phase and neural activity in the hippocampus (left) and amygdala (right). Across age groups, participants who perform better in the decision phase display greater activation during encoding in both of our regions of interest.

To model neural activity in the decision task, we fit three different GLMs. In the first, we included the following regressors of interest: 1) face trials on which the participant made an adaptive choice, 2) face trials on which the participant made a maladaptive choice, 3) onset for novel faces, 4) parametric effect of perceived amount given on all choice trials containing previously encountered faces. To examine neural evidence for participants’ tendency towards choosing faces regardless of value, we fit a second GLM that included the following regressors of interest: 1) onset of trials where the face was chosen, 2) onset of trials where the face was avoided, 3) parametric modulator for reaction time (z-scored). We then calculated the contrast between trials in which participants chose the face and trials in which participants avoided the face, the result of which is presented in Figure 3B and supplementary table 2. Finally, to highlight how perceived generosity may complicate this tendency towards choosing faces, we ran a third GLM that included the following regressors: 1) parametric modulator for reaction time (z-scored), 2) onset of low value high generosity faces, 3) onset of low value low generosity faces, 4) onset of high value high generosity faces, 5) onset of high value low generosity faces, 6) novel faces. The result presented in Figure 4C (and supplementary table 5) is the contrast between low value trials on which high versus low generosity faces were presented. Finally, because stimulus type is intermixed during the decision task, meaning that participants are making choices about both faces and houses in the same run, we also included regressors for house trials in all of our GLMs as follows: 1) parametric modulator for reaction time, 2) onset for adaptive house choices, 3) onset of maladaptive house choices, 4) onset for house trials with novel images. Since the focus of this paper is on social decision-making, however, we do not highlight those results here.

**Figure 3.**
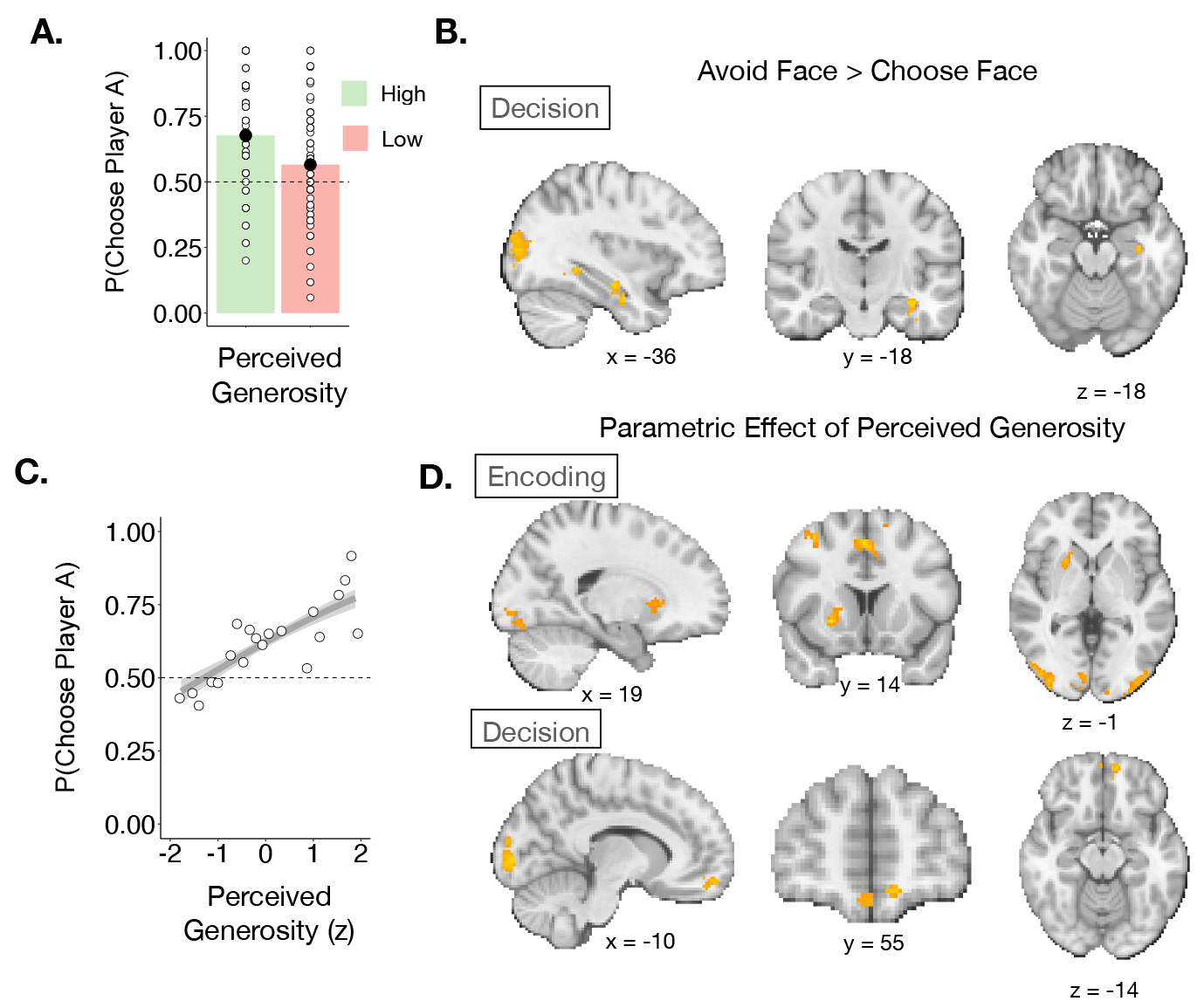
**(A)** Participants have a bias towards choosing the face over the outline, even when it does not look generous. **(B)** Across age groups, there is more hippocampal activation on trials where participants choose the outline over the face. **(C)** Perceived generosity (how much an independent group of raters thought each person would share in a hypothetical dictator game) predicts how often each face is chosen in the decision phase. **(D)** Neural signatures of perceived generosity at encoding and at decision.

**Figure 4.**
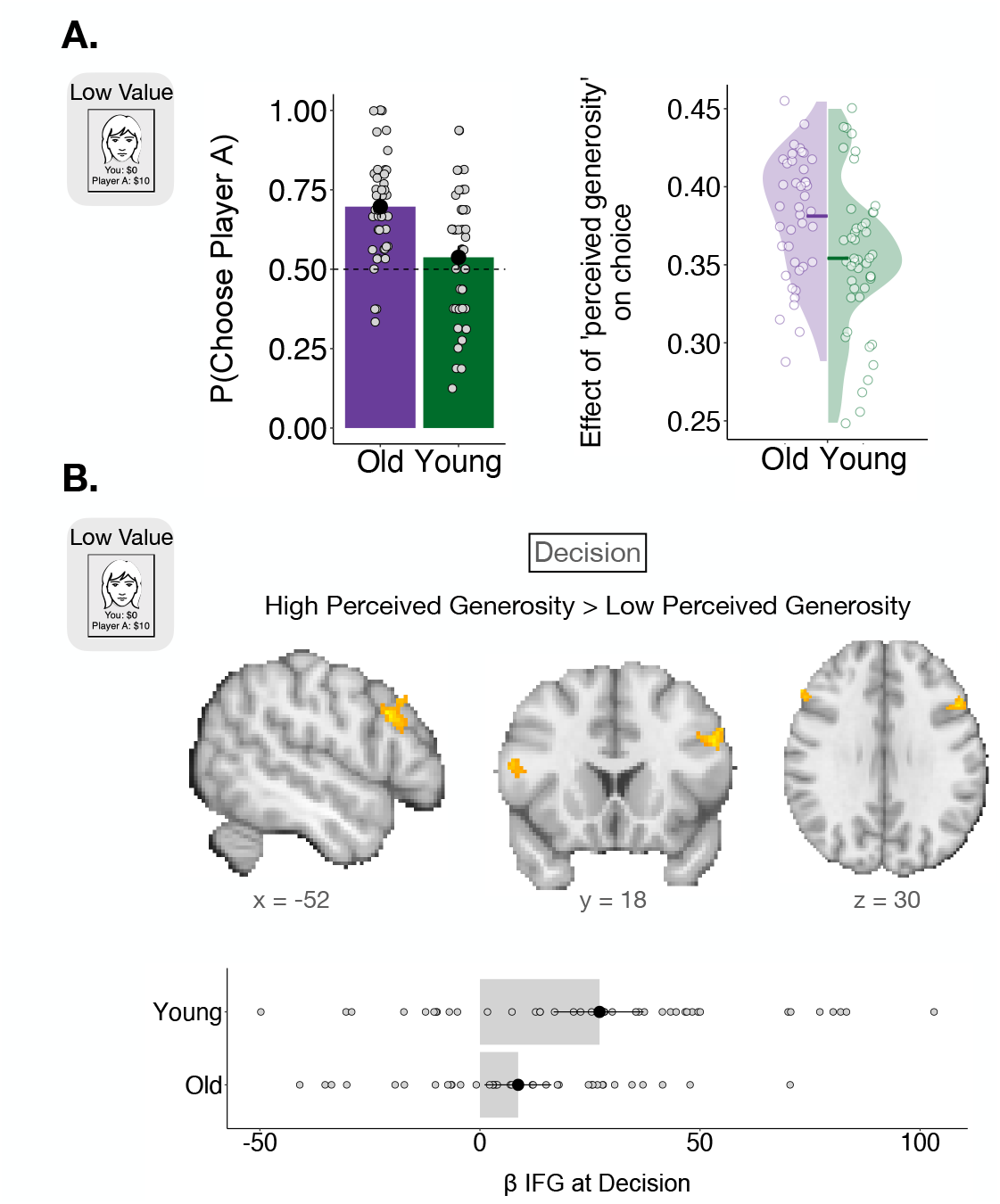
**(A)** *Left*. Proportion of choices in which the face was chosen, focusing on decision trials with low value faces. Older adults were significantly more likely to choose the face than younger adults were. *Right*. Individual coefficients for the effect of perceived generosity on choice during low-value trials. Older adults are more likely to be influenced by perceived generosity on trials when they should avoid the face. **(B)** There is more activation in the lateral prefrontal cortex (specifically, the IFG) on decision trials where the outcome associated with a face is at odds with how generous it looks. This increase in activation is greater for younger than older adults.

We were also interested in examining activity in ROIs that are relevant for memory-based decision making, such as the amygdala and the hippocampus. For the hippocampus and amygdala, we used ROIs derived from the Harvard-Oxford Atlas, applying a 25% threshold. This atlas contains a map of cortical and subcortical areas provided by the Harvard Center for Morphometric Analysis. We then used the *fslmeants* command is FSL to extract neural activation within the ROIs for the regressor of interest (namely, the onset of face encoding). We also conducted an exploratory ROI analysis using the functional ROI (fROI) that emerged from the low value high generosity > low value low generosity contrast. Within this fROI, we similarly used *fslmeants* to extract subject-specific betas that reflect the magnitude of neural activation in response to the contrast of interest. We then performed an independent-samples *t*-test on the extracted coefficients in order to test for reliable age differences.

## Results

We expected participants to leverage their memory for how much each person shared with them during the reward task to inform their choices during the decision phase. If participants remembered that a person had shared $5, they should choose to play with them again over a randomly drawn individual (and vice versa). Based on this logic, we can define an adaptive choice as a trial on which participants either avoided a low-value face or chose a high-value face. While young adults scored significantly above chance in the proportion of adaptive choices (t(44) = 2.19, p = 0.033), older adults did not (t(40) = -1.3, p = 0.17, Figure 1B). Similarly, whether a face was associated with high or low reward did not significantly predict choices for older adults (β_old_ = -0.16, p = 0.22), but it did for younger adults (β_young_ = 0.23, p = 0.037). In line with this differentiation, we found a significant interaction between age and the effect of reward on choice (β_Age*Reward_ = 0.41, p = 0.016).

One possible explanation for older adults’ poorer performance on this task is that they had difficulty remembering which faces they had previously seen. To assess whether impaired recognition memory could be at the root of the choice patterns we observe, we calculated a subject-specific d’ that captures how accurately participants were able to discriminate between old and new faces during the memory test. We found that both younger and older adults were above chance in their ability to recognize faces from the reward phase (t(44) = 15.09, p < 0.001, and t(39) = 10.6, p < 0.001, respectively), though younger adults were significantly more accurate than older adults (t(83) = -2.84, p = 0.005). In this task, however, recognition memory is not enough to support adaptive choice since it does not provide information about which of the faces was generous. For this reason, participants’ recognition memory score (d’) did not predict adaptive decision-making across age groups (r = 0.16, p = 0.15).

On top of recognizing the faces, participants also needed to remember the association between each face and how much they had shared to make an adaptive choice. Indeed, we found that associative memory predicts adaptive choice accuracy across the age range (β = 2.34, p < 0.001). Older adults, however, were not above chance in their ability to report whether a face from the reward task was associated with $0 or $5 (t(39) = 0.67, p = 0.50). They also had a “positivity bias” when remembering value, meaning that they were more likely to misremember a $0 face as a $5 face than the opposite (associative memory bias = -0.22, t(39) = -3.51, p = 0.0011). Young adults, on the other hand, could reliably recall the amount of money that was associated with each face (t(44) = 4.74, p < 0.001), and they did not display any bias in their memory for value (associative memory bias = 0.07, t(44) = 1.38, p = 0.17). Taken together, these findings suggests that age affects people’s ability to make decisions based on associative memory, at least in part because older adults have difficulty remembering how much each person shared with them.

Next, we were interested in assessing whether neural activity could help explain the decline in people’s ability to use associative memory in service of choice. In a whole-brain analysis looking at age-dependent differences at encoding, we found that in the 4.5 seconds during which faces were shown with their associated value, there was greater neural activation in the hippocampus and amygdala of younger than older adults (Figure 2A). Furthermore, when we extracted subject-specific estimates of neural activity at encoding from atlas-derived masks of the hippocampus and the amygdala, we found that activity in both brain regions at encoding positively predicted adaptive choice across the age range (hippocampus: r = 0.26, p = 0.007; amygdala: r = 0.23, p = 0.04; Figure 2B). This relationship suggests that activity in brain regions known to encode memory for valence (Phelps, 2004; Phelps et al., 1997) may support value-based decisions that rely on previous experience. Thus, age-related changes in neural activity in these brain regions can help explain the behavioral differences we observe.

Although participants should use value memory to guide their decisions, there are other features of the experimental stimuli that could bias choice. Indeed, even when participants had correct memory for value, their average adaptive choice accuracy was only 65%. We hypothesized that this might be due to the inherently social nature of the experiment, which could affect performance in at least two ways: participants might be generally biased towards choosing faces rather than silhouettes, and participants might make choices on the basis of how generous a face looked rather than episodic memory. In Lempert et al. (2022), the authors found that, indeed, older adults tended to be especially influenced by these features, weighing perceived generosity more strongly than younger adults and approaching more faces overall. In this sample, we expected to find similar behavioral patterns, coupled with neural evidence of their influence.

On one hand, we found that participants of all ages demonstrated a “face bias” such that, in general, they tended to choose the face over the silhouette (Figure 3A). Indeed, across all levels of perceived generosity and associated value, participants chose the face on 62% of trials, which is significantly above chance (one sample *t*-test: t(85) = 6.4, p < 0.001). Given this bias towards choosing faces, we hypothesized that when participants chose the silhouette, they might be relying more strongly on associative memory. To assess this claim, we compared neural activity on choice trials when participants chose the face versus when they chose the silhouette. As predicted, we saw greater activity in the anterior hippocampus when participants did not choose the face, regardless of age (Figure 3). Since the hippocampus is known to support associative memory, this finding suggests that participants’ bias towards choosing faces can sometimes be overridden by intact memory for value.

Beyond this face bias, participants’ choices are also likely to be biased by how generous each person looks. Across both age groups, we found a significant effect of perceived generosity on choice (Figure 3C). People were more likely to choose to play with faces that look more generous, as established by a group of independent raters (β = 0.42, p < 0.001; see Methods for more details about the rating procedure), and age did not modulate how much people leveraged these irrelevant facial attributes to guide behavior (β = -0.04, p = 0.66). Neurally, this sensitivity to facial features was evident during both encoding and retrieval. Indeed, when participants first saw the images, we found that activity in the striatum parametrically tracked how generous a face looked (Figure 3D). This striatal effect was present regardless of age, echoing the behavioral finding. Later, at the time of decision, activity in the ventromedial prefrontal cortex (vmPFC) was modulated by perceived generosity, suggesting the unfolding of an evaluative process that takes this feature into account (Figure 3D).

So far, we have identified behavioral and neural correlates for three different mechanisms that help determine people’s choices – value memory, a bias towards choosing faces, and perceived generosity. Only the neural correlates of value memory encoding appeared to change with age; we found neural and behavioral evidence for sensitivity to facial features across the age range. Still, older adults’ worse performance cannot exclusively be explained by worse value memory: even on trials where they reported intact memory, their choices were less value-driven than younger adults’ (β_Age*Reward_ = 0.84, p = 0.014). The question remains, then, of how to explain older adults’ reduced ability to avoid low value faces, beyond the memory impairments we observe.

The answer to this question emerges if we consider choice behavior as a product of multiple competing influences (memory, face bias, and perceived generosity) that are operating simultaneously. Sometimes, these different factors all lead to the same decision – for instance, when a high-value generous-looking face is presented. Other times, the tendency to choose faces and to do so according to how generous they look may be at odds with how much the person shared in the reward task. Under these conditions, participants should prioritize their memory for how much someone actually gave them, suppressing their other biases. We found that this process of inhibiting the influence of irrelevant information was impaired in older adults, both in terms of their tendency towards choosing the face and the influence of perceived generosity. Indeed, focusing on decision trials that most evoke this tension – those in which a low-value face is presented – we found that older adults were significantly more likely to erroneously choose the face (t(85) = -4.05, p < 0.001, Figure 4A), even when they remembered later that it was unrewarded (t(78) = 2.77, p = 0.0068). On these low-value trials, older adults were also significantly more likely to base their decisions on perceived generosity (*t*-test on random effects: t(84) = 2.8, p = 0.005; Figure 4B) rather than episodic memory. Taken together, these results suggest that while both younger and older adults were sensitive to the same biases, younger adults were better able to override them in favor of associative memory when it was necessary to do so.

Based on this finding, we might expect to see age-related differences in neural activity when participants have to make choices that most rely on suppressing irrelevant information – namely, when a low-value face looks generous. Indeed, these are trials in which participants need to avoid the face, going against both their bias towards choosing faces and their tendency to choose according to perceived generosity. On these trials, we found increased activation in the inferior frontal gyrus (IFG) and lateral prefrontal cortex (lPFC) more generally (Figure 4C). Importantly, the increase in dlPFC activity was significantly greater in the brains of younger than older adults (t(72) = 2.77, p = 0.0071). Given that the dlPFC has been linked to inhibiting the influence of goal-irrelevant information (Miller & Cohen, 2001; Thompson-Schill et al., 2005), this finding provides neural evidence for the notion that younger adults are better able to suppress the influence of irrelevant features on choice. In sum, both memory for valence and the ability to adaptively deploy it in service of choice contribute to successful performance on this task. We find neural and behavioral evidence to suggest that aging affects both components of this process.

## Discussion

In this study, we further our understanding of how social decisions are affected by aging by identifying the neural correlates of (1) reduced memory for people’s past behavior and (2) reduced inhibition of irrelevant social stereotypes. We see decreased activation in older adults’ MTL at encoding, which predicts less adaptive decision-making later. Furthermore, while both younger and older adults show activity in value-responsive brain regions that tracks with perceived generosity, younger adults are better able to adaptively suppress the influence of this factor that is not goal-relevant, an inhibitory process accompanied by increased activity in dorsolateral regions of the prefrontal cortex in younger adults.

While providing new insights into how age-related changes in brain function lead to systematic changes in social decision-making, these findings are also consistent with previous work. First, the notion that the hippocampus contributes to the formation and maintenance of episodic memory is well-established. Patients with hippocampal lesions cannot form new memories of specific events (Scoville & Milner, 1957; Zola-Morgan & Squire, 1986), optogenetic interventions to the mouse hippocampus can reactivate or inhibit specific engrams (Liu et al., 2014; Ramirez et al., 2014), and human fMRI studies regularly implicate the hippocampus at both encoding and retrieval (Chadwick et al., 2010; Rugg & Vilberg, 2013). Although the exact mechanism by which the hippocampus supports episodic memory is not fully agreed upon, its role is especially evident when different elements of an experience must be bound together (Maguire et al., 2016; Mullally et al., 2012; Roberts et al., 2018). In this paradigm, the memory that is required for successful performance is inherently associative, since participants need to remember the link between faces and how much they gave. For this reason, our finding that hippocampal activity predicts performance is well-aligned with existing research. In fact, a recent study using a similar paradigm in a cohort of younger adults identified the time-course of hippocampal activation as the primary determinant of choice (FeldmanHall et al., 2021). While FeldmanHall et al. (2021) focused on neural activity at decision time, however, we highlight the importance of hippocampal activity at encoding and show that it is correlated with adaptive choice.

Importantly, we also identify the amygdala as playing an important role in value-based choice. In line with this finding, the amygdala can support memory for valence (Cahill, 2000; Kensinger & Corkin, 2004; Phelps et al., 1997b), helping people preferentially approach stimuli that yield positive outcomes (Cahill, 2000; Kensinger & Corkin, 2004; Phelps et al., 1997b). Furthermore, part of the amygdala’s role in supporting memory for emotional cues may be driven specifically by its interactions with the hippocampus (Phelps, 2004b; Richter-Levin & Akirav, 2000). In rats, for instance, stimulating the basolateral amygdala reinforces long-term potentiation in the hippocampus, which is at the root of long-term memory (Frey et al., 2001; Wittmann et al., 2008). In the present study, we find that increased activation in both the hippocampus and the amygdala during encoding tracks with appropriately using valence to guide later choices, supporting the idea that these regions work together to support long-term memory for item valence. If the hippocampus and the amygdala facilitate the encoding of valence, it follows that reduced activity in these brain regions would lead to suboptimal choice. In our sample, the neural and behavioral findings in older adults both corroborate this prediction. This result is in line with other research that has identified changes to the structure and functioning of amygdala and hippocampus with age. Indeed, both the hippocampus and the amygdala display atrophy as people age (Aghamohammadi-Sereshki et al., 2019; Buckner et al., 2006; Kurth et al., 2019; Lister & Barnes, 2009; Malykhin et al., 2008; Raz et al., 2005), and the degree of atrophy is related to cognitive symptoms of preclinical Alzheimer’s Disease (Wolk et al., 2017). In the hippocampus, these changes have been explicitly linked to impairments in episodic memory and the precision with which participants can recall images (Stark et al., 2019; Yassa et al., 2011). Furthermore, older adults exhibit decreased functional connectivity between the hippocampus and amygdala, which is linked to worse subsequent memory for negative stimuli (St. Jacques et al., 2009). Prior research has largely focused on the relationship between neural functioning in these regions and memory. Here, we extend these past findings by showing how changes in neural functioning of these areas can also have consequences for decision making.

Though impaired associative memory undoubtedly contributes to worse decision-making in older adults, we also know that participants do not rely exclusively on memory to make their decisions. They also make inferences about others’ personality traits based on their facial features (Todorov et al., 2008, 2009), and leverage this information to decide whom to approach. In our sample, this evaluative process is subserved by the striatum and vmPFC, both of which are central hubs of the brain’s valuation network (Bartra et al., 2013; Henri-Bhargava et al., 2012; Kang Souther et al., 2023; Levy & Glimcher, 2012; Schultz et al., 1997). Interestingly, while the striatum is sensitive to facial features at encoding, activity in the vmPFC is modulated by perceived generosity at choice. This dissociation suggests that the striatum is more strongly implicated during learning, whereas the vmPFC is more strongly recruited at decision time. This is in line with previous evidence that highlights the relative activation of the striatum and vmPFC during learning and choice (Bartra et al., 2013; Jocham et al., 2011). In the realm of social decision making specifically, the vmPFC is also known to track with inferences of other’s generosity (Cooper et al., 2010) and to play a role in facial emotion recognition (Heberlein et al., 2008; Hiser & Koenigs, 2018; Tsuchida & Fellows, 2012). The orbitofrontal cortex, which consists of the ventral portion of the vmPFC, has also been found to encode stereotypes in the context of social decision-making (Kobayashi et al., 2022; Stolier & Freeman, 2016). In general, the vmPFC may be broadly involved in inferring valence in service of decisions, including on the basis of irrelevant facial attributes.

While both younger and older adults are more likely to approach familiar people regardless of their past behavior, we found that this bias is stronger in older adults. One possibility is that this bias arises from an aversion to uncertainty, since the alternative option is to interact with a randomly chosen past participant. Indeed, older adults are sometimes more uncertainty-averse than younger adults, particularly when choices are risky (Frank & Seaman, 2023; Rolison et al., 2014; Tymula et al., 2013) – though evidence to the contrary also exists (Mata et al., 2011). Another explanation for older adults’ tendency to choose familiar individuals is a positivity bias (Carstensen & Mikels, 2005; Mather & Carstensen, 2005), a phenomenon by which older adults are more likely to disregard negatively-valenced information and focus their attention on positive outcomes. Lempert et al. (2022) found evidence for a positivity bias, in that the older adults were more biased to remember faces as generous. Even when they did accurately remember that an individual was not generous, older adults were more likely to choose to interact with them anyway. We replicate both results in our sample, suggesting that older adults might be generally optimistic about or unbothered by players who proved themselves to not be generous. This kind of reappraisal, coupled with a potential aversion to uncertainty, may help explain the increased familiarity bias that emerges with age.

Finally, although we find no age differences in sensitivity to perceived generosity (in either our neural or behavioral data), older adults are more likely to choose according to those perceptions when they should not. In these moments of tension between episodic memory and judgments based on stereotypes, age modulates which source will be more influential on choice. In the non-social domain, Lalla et al. (2022) have reported similar age-dependent changes in how episodic versus semantic information is used to guide choice. They found that when participants were asked to choose between two objects, older adults were more likely to base their choices off the items’ real-world value, as opposed to the value that was assigned during the task. Younger adults were better able to suppress task-irrelevant semantic information, which allowed them to make more adaptive choices overall. In part, these age differences may be driven by decreased inhibitory control in aging, which makes it difficult to override biases that would lead them to suboptimal choices. Indeed, the inhibitory deficit hypothesis posits that age-related declines in performance across memory and decision-making tasks result from older adults’ difficulties in suppressing irrelevant information (Hasher et al., 1999; Hasher & Zacks, 1988). From a memory perspective, this leads to more cluttered memory representations, including the overrepresentation of irrelevant semantic knowledge (Healey et al., 2013). Successful arbitration between competing sources of information may rely on neural substrates like the dlPFC that are known to support cognitive control (Botvinick & Cohen, 2014) – including the ability to select relevant information from memory (Chrysikou et al., 2014; Thompson-Schill et al., 2005) and weight the different dimensions of available choice options (Hutcherson et al., 2012). Together, these findings suggest that reduced dlPFC activation in older adults reflects a disruption in the process by which stimulus features are recruited in service of choice.

In this study, we used fMRI data to better understand how and why older adults are biased towards approaching people it is economically disadvantageous for them to trust. Our results pave the way for future research in several ways. For instance, we find that decreased MTL activation at encoding predicts suboptimal choices. One question that remains is whether older adults rely more on perceived generosity because they cannot remember value, or whether they do not encode value in the first place because it is not likely to significantly influence their decisions. To address this question, it may be helpful to probe participants’ metacognitive awareness of which factors led to their choices. Considering older adults’ metacognition could also help us uncover other novel explanations for their behavior. For instance, socioemotional selectivity theory (Carstensen, 2021) posits that as people begin to view their time on earth as more limited, they will re-orient their social preferences to maximize their well-being. It may be that assuming the best in others is a component of this adaptive reappraisal, leading to greater life satisfaction overall. Another interesting metacognitive question is whether older adults are more likely to believe that social stereotypes, like perceived generosity, really do predict behavior. If so, this could also call into question whether the patterns of behavior we observe are truly irrational, or rather driven by (incorrect) explicit beliefs. Future work would also be helpful for establishing the generality of the results observed here, and to leverage this mechanistic knowledge to help train older adults to overcome these biases. Despite any limitations in applying new episodic information, older adults still have valuable semantic knowledge to draw upon in making decisions. Thus, it is important to understand both the strengths and the limitations of older adult decision makers to best position them, and society, to be strategically effective.

## Supplementary Tables

Note: Brain regions defined probabilistically using the Harvard-Oxford Atlas

**Figure 2A:**
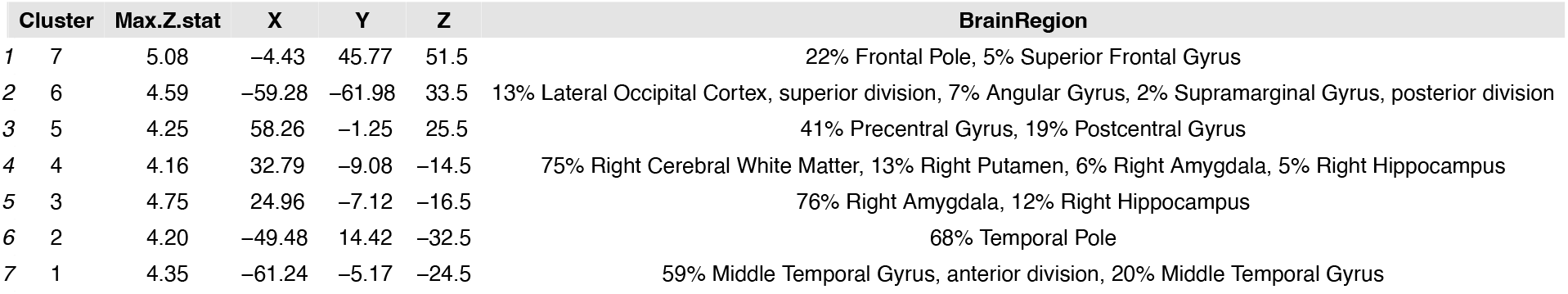
whole-brain, young > old significant peaks

**Figure 3A:**
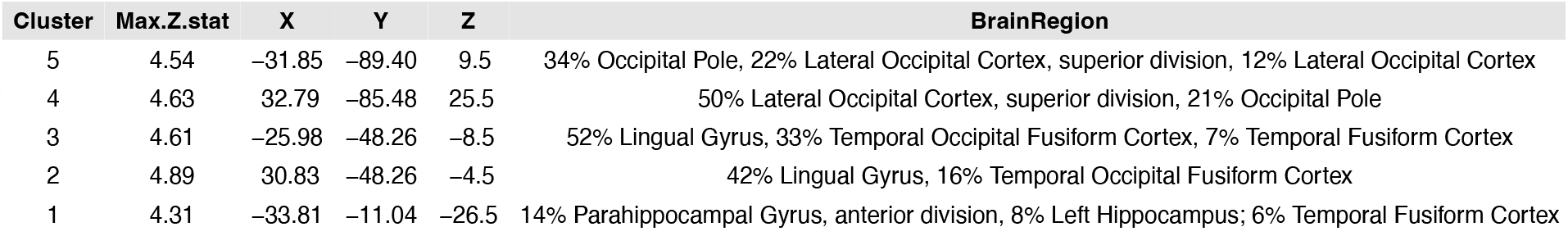
whole-brain, avoid > choose significant peaks

**Figure 3D:**
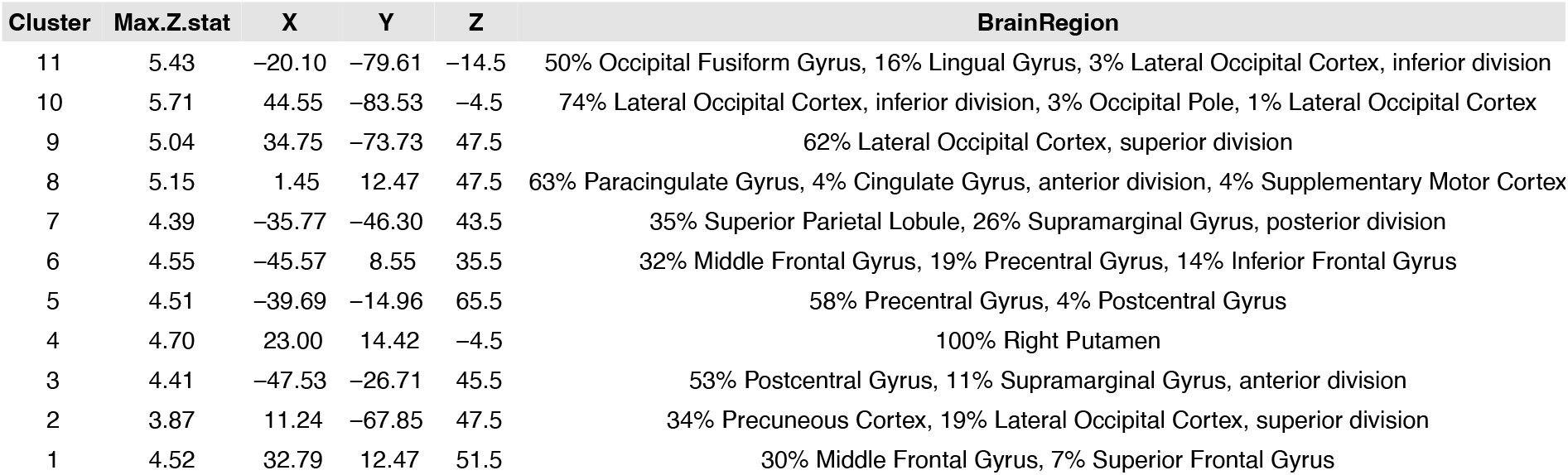
whole-brain parametric effect of perceived generosity (encoding)

**Figure 3D:**
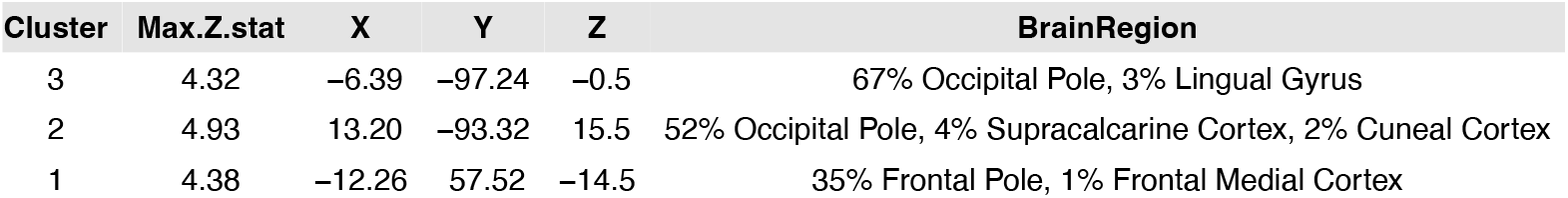
whole-brain parametric effect of perceived generosity (decision)

**Figure 4B:**
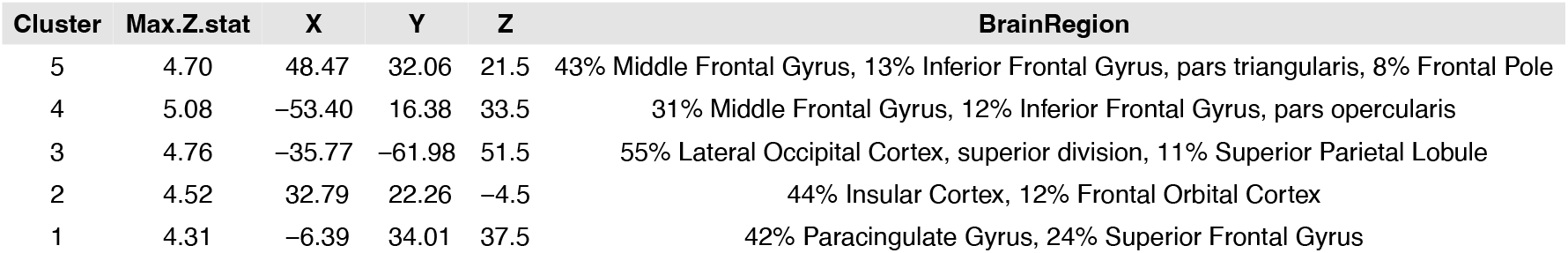
whole-brain, high perceived generosity > low perceived generosity

